# Early seral vegetation communities increase insect abundance and diversity in a semiarid natural gas field during early and late growing season

**DOI:** 10.1101/2022.06.28.494893

**Authors:** Michael F. Curran, Joshua R. Sorenson, Zoe A. Craft, Taylor M. Crow, Timothy J. Robinson, Peter D. Stahl

## Abstract

Insects are critical components of terrestrial ecosystems and are often considered ecosystem engineers. Due to the vast amount of ecosystem services they provide, because statistically valid samples can be captured in short durations, and because they respond rapidly to environmental change, insects have been used as indicators of restoration success and ecosystem functionality. In Wyoming (USA), thousands of acres of land surface has been disturbed to extract natural resources. While traditional reclamation practices of these lands focused on site stabilization and weed control, more recent efforts have been made to restore ecosystem services. It has been suggested that a spatial and temporal mosaic of flowering species will benefit insect populations. In this study, we compared early seral reclamation sites (i.e., well pads undergoing interim reclamation) to reference areas at two points within a growing season. We found reference ecosystems were devoid of forb species, while one year old reclaimed sites contained late-season blooming Rocky Mountain beeplant (*Cleome serrulata*) and three-four year old well pads contained early-season blooming perennial forb species, mainly western yarrow (*Achillea millefolium*). We compared insect abundance and family richness on 6 well pads with early season perennial forbs and 6 well pads with the late season annual forb, Rocky Mountain beeplant to insect communities on adjacent reference areas. A total of 237 insects were found on early season reclaimed sites compared to 84 on reference sites, while 858 insects were found on late season reclaimed sites compared to 40 on reference sites. Insect abundance and family richness was significantly higher on reclaimed well pads compared to reference areas at both points in the growing season. We also found interesting differences in abundance at family levels.

## Introduction

Insects, the most abundant and diverse group of animals in the world, provide numerous ecosystem services (Losey & Vaughan 2006, Noriega et al. 2018). Insects are often considered ecosystem engineers because they impact soil properties (Lavelle et al. 1997), influence nutrient cycling (Belovsky & Slade 2000), serve as a food source to higher trophic levels (Tallamy 2009), and provide critical pollination services allowing plants to successfully reproduce (Archer & Pyke 1991, Dixon 2009, Cusser & Goodell 2013). Since they provide more ecosystem services than other wildlife, respond rapidly to environmental change, and because statistically valid samples can be captured in a short duration, insects have been used as indicators of restoration success (e.g., Kimberling et al. 2001, Longcore 2003, Rohde et al. 2019). While traditional focus of insects in ecological restoration projects has been towards pollinators in agriculture systems (Menz et al. 2010), more recently they are becoming more common in non-crop studies (e.g., Cusser & Goodell 2013, Tonietto & Larkin 2018, Rohde et al. 2019, Curran et al. 2022).

In the western United States, extraction activities related to energy development have resulted in hundreds of thousands of acres of land surface disturbance. In Wyoming, oil and gas operating companies are required to reclaim surface disturbances after extraction activities take place while complying with multiple regulatory criteria (Curran et al. 2013, 2019a). These criteria have typically focused on establishing vegetation deemed suitable by regulatory agencies with a heavy emphasis on controlling erosion (Curran & Stahl 2015). More recently, focus has shifted from strictly land reclamation towards restoring disturbed landscapes to functional ecosystems, providing suitable wildlife habitat (Stahl & Curran 2017). Insects and other arthropods, as wildlife which serve as ecosystem engineers and play significant roles as bottom-up drivers of trophic interactions (Grodsky et al. 2018), have been used as indicators to determine success of ecosystem restoration projects (Longcore 2003), but have rarely been studied in oil and natural gas fields (Sylvain et al. 2019; Curran et al. 2022).

The first study to examine how insects respond to ecosystem restoration efforts associated with natural gas development in a sagebrush-steppe ecosystem suggests these efforts increase insect abundance and diversity (Curran et al. 2022). The mass flowering hypothesis, which suggests dense patches of flowering plants increase pollinator abundance in areas where the surrounding landscape is void of diverse floral vegetation (Westphal et al. 2003, Holzschuh et al. 2013, Todd et al. 2016), was tested in that study. A limitation of the study, which occurred in the Pinedale Anticline natural gas field, is it only examined one flowering species, Rocky Mountain bee plant (*Cleome serrulata;* hereafter RMBP) at one point in the growing season. This issue is found among studies which have tested the mass flowering hypothesis in agricultural systems. Research suggests having a diverse spatial and temporal mosaic of flowering plants which bloom at different periods throughout the growing season is an effective strategy for insect and pollinator conservation (e.g., Samways et al. 2020).

In the Jonah Infill natural gas field (hereafter Jonah Field), well pads are constructed by stripping topsoil and vegetation and stockpiling it to allow for drilling equipment to operate on level surface. Typically, 70-80% of initial land surface disturbance has soil respread and seeded the same year construction equipment is removed from the site. The most commonly used seed mix in the Jonah Field contains 21 native plant species (6 perennial grass species, 4 shrub species, 10 perennial forb species and 1 annual forb; Supplemental Table 1). The annual forb used in the seed mix is RMBP. It is common for the annual forb, Rocky Mountain bee plant, which grows well in disturbed soil (Cane 2008) to be the most prolific species in terms of establishment and flowering in the first year after reclamation. However, it tends to become less dominant within two-three years as perennial forbs, shrubs and grasses establish (Curran et al. 2019b). While RMBP flowers later in the season, many perennial forbs in the seed mix flower earlier in the season. The objective of this study was to compare insect communities on natural gas well pads in the Jonah Infill natural gas field (Sublette County, Wyoming, USA) undergoing restoration activities at two points in the growing season to determine if insect communities were similar at different parts of the growing season while different floral communities were in bloom. We examined vegetation and insect communities on six, three year old well pads in the early portion of the growing season (June 22, 2017) which contain diverse perennial forbs which all bloom earlier in the season than Rocky Mountain bee plant on nearby well pads that have been seeded that year. Later in the season (July 27, 2017), when the flowers on three year old sites were mostly done blooming we examined vegetation and insect communities on six, one year old well pads with prolific flowering from Rocky Mountain bee plant (*Cleome serrulata*). In both instances vegetation and insect communities on adjacent reference sites were examined. Since the adjacent reference sites consist primarily of decadent Wyoming big sagebrush (*Artemisia tridenta* spp *Wyomingensis*), with little understory and very few or no forbs, we hypothesized insect communities on reclaimed well pads would have more abundant and diverse insect communities than adjacent reference areas. We also sought to determine if differences in insect communities existed between early and late season well pads and reference sites.

## Methods

### Study Area

A total of 12 well pads undergoing interim reclamation and their adjacent reference areas were monitored in the Jonah Infill natural gas field in Sublette County, WY, USA). This field is predominantly Federal land and is regulated by the Jonah Interagency Office (JIO – consisting of Pinedale BLM Field Office, Wyoming Department of Agriculture, WYDEQ, and Wyoming Game and Fish Department). Well pads in this study averaged 5.3 acres of initial surface disturbance with 70-80% of that area undergoing interim reclamation (i.e., soil respread and seeded on non-active portions of the well pad).

All well pads were located in the Stud Horse Butte (SHB) section of this gas field. All sites were located in ecological site R034AC122WY. This ecological site description is dominated by loamy soils. The historic climax plant community is described as ‘big sagebrush/bunchgrass’, though the state and transition model for the site suggests continuous high intensity early season grazing may change the historic climax plant community to ‘big sagebrush/bareground’. Grazing records were unavailable, though it has been documented the area has been grazed primarily in the Spring since the early 1870s (Sommers 1994). This site description averages between 30 and 70 frost free days per year, with mean annual temperatures between 1.1^°^C-3.3^°^C. The average annual precipitation, according to the ecological site description, is 22.9 cm – 30.5 cm. The elevation of the area ranges from 2,136 m – 2,219 m.

All well pads were seeded with seed mix ‘B1’ (Supplemental Table 1). The mix contains four native shrub species, 10 native perennial forb species, one annual forb species, and six native perennial grass species. Six of the well pads were 3-4 years old and consisted mainly of native perennial forbs, grasses and shrubs. These sites were sampled on June 22, 2017 while perennial forb species were in bloom. The other six sites in this study were in their first year of reclamation and were sampled on July 27, 2017 when the native annual forb, *Cleome serrulata* (RMBP), was in bloom. The primary vegetation on these locations was Rocky Mountain bee plant (*Cleome serrulata*).

### Vegetation Sampling

Well pads which consisted of early season blooming flowers were sampled on 22 June 2017, while later season flowers were sampled on 27 July 2017. Two forty meter transects were sampled at 5 m and 10 m away from the edge of the well pad, both in the reference area and on the well pad. A 0.5m^2^ image was taken at the start of each transect and at 5m increments along the transect, resulting in 9 images per transect, or 18 for each well pad and associated reference area. The images were taken using a free-hand nadir technique (Cagney et al. 2011) and had ground sample distances of ∼0.4mm which is similar to previous studies examining vegetation on reclaimed well pads (Curran et al. 2019a, 2020). A 7×7 grid was placed in each image in ‘SamplePoint’ (Booth et al. 2006). This resulted in a total of 49 pixels being classified per image, or 441 pixels per transect (882 pixels per each well pad and associated reference area). Pixels were classified to the species-specific level for shrubs and forbs, while grasses were classified as rhizomatous or bunch grass. Bareground, rock, and litter were also classified in pixel analysis.

### Insect Sampling

Insect sampling was conducted on the same days as vegetation sampling. Two runs of forty sweeps were taken on each reclaimed well pad and adjacent reference location transect using a 38 cm diameter sweep net with a 50 cm handle. All sampling was conducted between the hours of 0730 and 1600. Sampled insects were transferred from the sweep net to a Zip-Lock^®^ bag and placed in a cooler located in a research vehicle parked in the long-term disturbed portion of the well pad, >50 m away from transect locations. In all instances, two individuals walked along the transect, with the lead individual conducting the sweep net collection and the follower taking freehand nadir images as described above. A five-minute wait period was conducted by both individuals between sampling transects to avoid ‘flushing’ effects on insects (Wenninger and Inouye 2008).

Upon return to the University of Wyoming (Laramie, WY, USA) from the field, samples were placed in a freezer for laboratory identification at a later date. Insects were Identified to family level under a microscope with aid from National Audubon Society field guide (Milne and Milne 2011).

### Statistical Analysis

Spreadsheet reports generated from SamplePoint analysis were used to to classify vegetation cover into bare ground, litter, and vegetation by species. Vegetation was classified into the functional groups forb, grass, or shrub. Wilcoxon signed-rank tests on paired differences were used to compare percent cover for bare ground, litter, forb, grass, and shrub between reclaimed sites and reference sites for each sampling period (Figure 1). Since these sampling events took place at different time points, early season sites and late season sites were compared separately.

**Figure 1.**
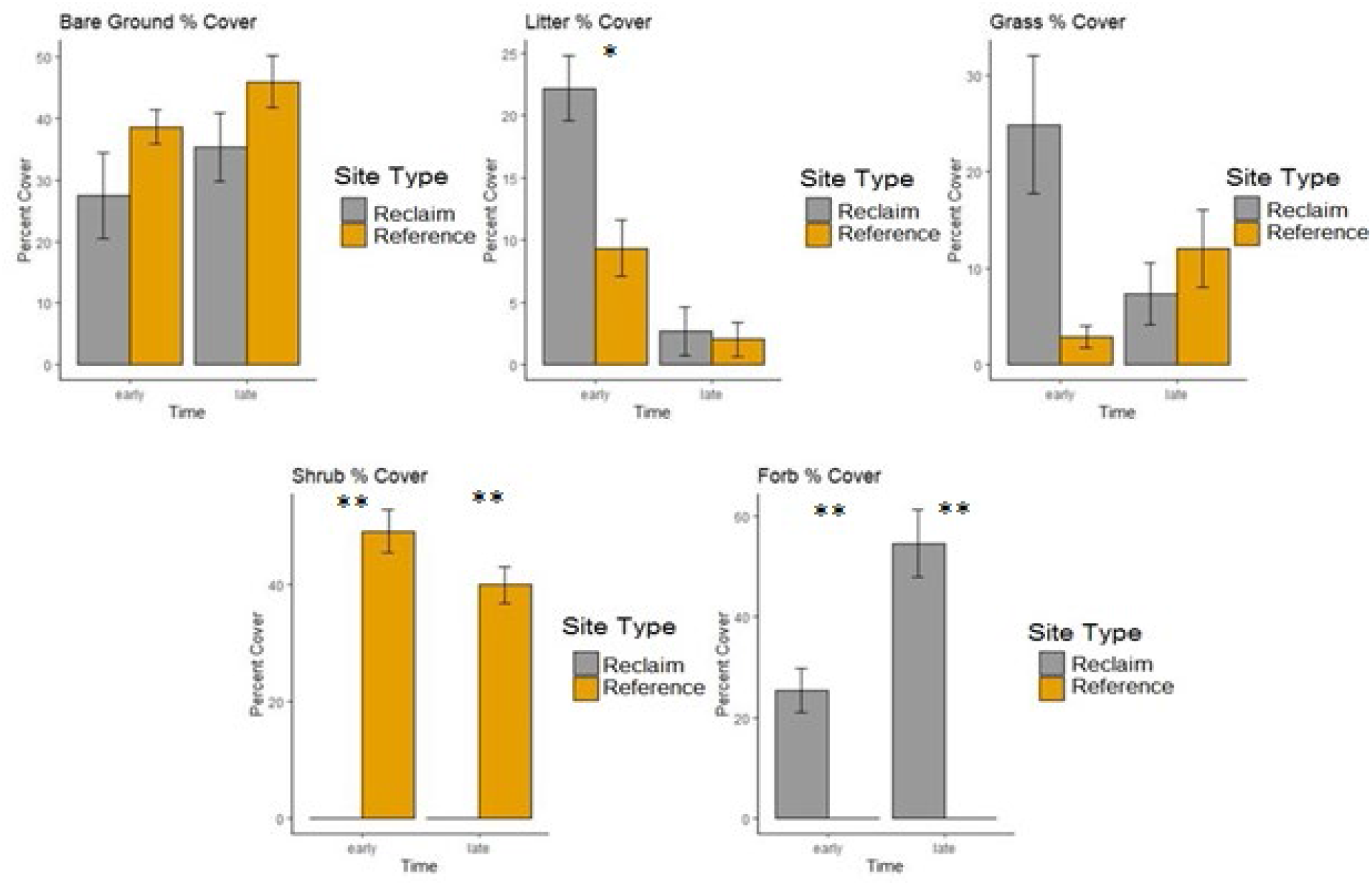
Bar graphs showing percent cover of (a) bare ground, (b) litter, (c) grass, (d) shrubs, and (e) forbs on reclaimed and reference sites which were monitored in early season and late season. *=p-value<0.05, **=p-value<0.001

For insect community analysis, sites were separated sites into the following groups before comparison: (1) 3-4 year old well pads monitored in early season, (2) reference sites adjacent to early season sites, (3) one year old well pads monitored later in season, and (4) reference sites adjacent to later season sites. Wilcoxon signed-rank tests were used to make comparisons between insects on reference vs. reclaimed sites in early and late season as well as comparisons between insects within reclaimed and reference sites in early and late season. Wilcoxon tests were also used to compare individual families within each of the original 4 contrasts.

## Results

### Vegetation Sampling

Vegetation communities on transect locations on reclaimed well pads differed from reference areas (Figure 1). Percent forb coverage was significantly greater on reclaimed well pads than reference areas (p<0.001) where no forbs existed in either the early or late season collection period (Figure 1). While 100% of forb coverage on reclaimed sites in the later growing season was RMBP (*Cleome serrulata*), upwards of 95% of forb coverage on early season sites was western yarrow (*Achillea millefolium*). Other forbs on early reclaimed sites included blue flax (*Linus lewisii*), Palmer’s penstemon (*Penstemon palmeri*), littleflower penstemon (*Penstemon procerus*), scarlet globemallow (*Sphaeralcea munroana*), and evening primrose (*Oenothera pallida*). Percent shrub coverage was significantly greater on reference areas than reclaimed sites (p<0.001) where shrubs were not present within transect locations (Figure 1). All shrubs in the reference areas were Wyoming big sagebrush (*Artemisia tridentata* spp. w*yomingensis*). Percent litter coverage was significantly higher on reclaimed sites than reference areas in the early season (p=0.0355). No other significant differences existed between percent coverage categories (Figure 1). Although it appears as percent grass cover was significantly greater on early season reclaimed sites than reference areas, a p-value=0.0591 suggests the sites were not significantly different under p=0.05 criteria.

### Insect Sampling

Insect abundance was significantly different among site types with more insects found on reclaimed areas than adjacent reference areas (Figure 2). The mean amount of insects found on early season well pads was 39.5 compared to 14 on adjacent reference sites, meanwhile the mean amount of insects on late season well pads was 143 compared to 6.7 on their reference sites. A total of 237 insects were found on early season reclaimed sites with 84 on the reference sites, whereas 858 insects were found on late season sites and 40 were found on the reference sites.

**Figure 2.**
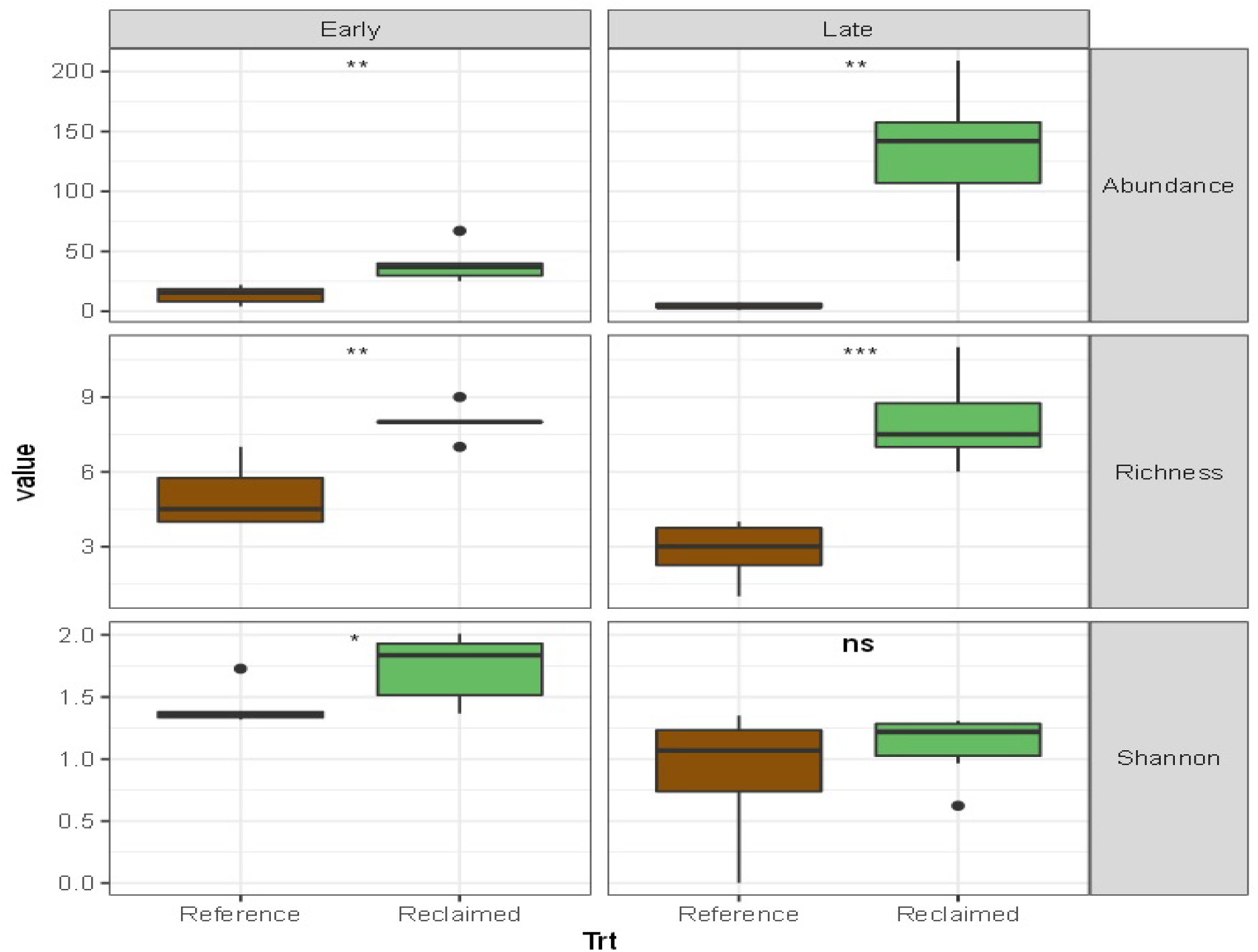
A graphical representation of abundance, richness and Shannon’s diversity index comparing reference vs reclaimed within early season and within late season. *=p-value <0.05, **=p-value<0.01, ***=p-value<0.001.

Insect richness was significantly different among site types with reclaimed areas having greater family richness than reference areas (Figure 2). A total of 24 insect families were identified in this study. A total of 18 insect families were found in the early season collection and 15 were found in the late season collection (Figures 2 and 3). Of these, nine families were common to both collections (Braconidae, Chalcidlidae, Cercopidae, Coccinellidae, Chrysomelidae, Formicidae, Halictidae, Lygaeidae, and Sarcophagidae). No insect families were significantly greater in reference areas than reclaimed areas during either early or late season collections (Figure 3). In early season, Phoridae and Miridae were significantly more abundant on reclaimed sites than reference areas (p<0.05; Figure 3). In late season, Chrysomelidae, Lygaeidae, Bombyliidae, Halictidae, and Formicidae were significantly more abundant on reclaimed well pads compared to reference areas (p<0.05; Figure 3). Chrysomelidae, Lygaeidae, and Bombyliidae were significantly more abundant on late season reclaimed sites than early sites (p<0.05; Figure 3), while the opposite was true for Phoridae, Miridae, and Cicadellidae (p<0.05; Figure 3). Cicadellidae were significantly higher in early season reference areas than late season reference areas (p<0.05; Figure 3), while no other insect families showed significant differences between reference sites at early and late collection periods.

**Figure 3.**
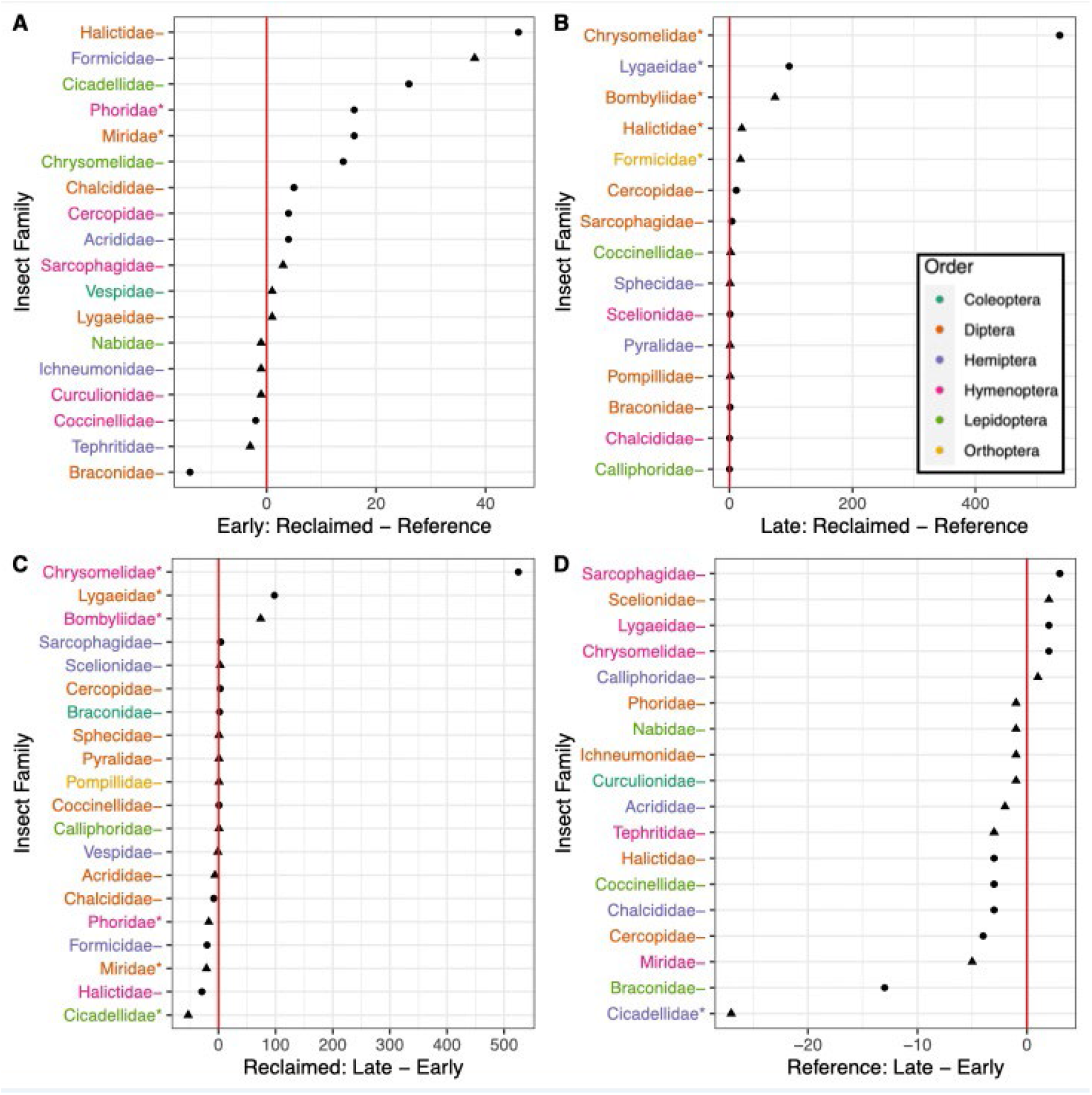
Total family level difference within each of 4 contrasts. A) Reclaimed minus Reference within Early season. B) Reclaimed minus Reference within late season. C) Late minus early within reclaimed and D) Late minus early within reference. Triangle point represents families that were only found in one of the two conditions. Families with ‘*’ next to them are those which had significant differences between site type with p<0.05.

## Discussion

As predicted, we found more insect abundance and diversity on restoration sites than the surrounding reference system. While we cannot rule out climatic factors as a potential cause for there being fewer insects in the first collection, previous research has shown yarrow, penstemon and blue flax, which were the main flowering species in bloom during the first insect collection, attract less wild bees than other native perennial forbs which bloom around the same time (Cane and Love 2016). However, it is not surprising that the well pads with early blooming flowers contained greater insect abundance and diversity than the adjacent reference system which had zero flowering species and were predominately composed of decadent sagebrush and bareground. Our results from late season sampling corroborate the previous study to examine insects on well pads containing Rocky Mountain bee plant and adjacent reference sites (Curran et al. 2022).

While we were not able to determine if insects were moving among well pads at either sampling time of our study, it is possible that providing beneficial resources (i.e., flowers) in a patchy matrix throughout the existing landscape could improve insect movement, reproduction, and increase overall abundance and diversity of insects in the area (Cane 2001). If insects are, in fact, moving among well pads, they may be delivering benefits to plants in the reference areas by increasing genetic diversity through cross-pollination and may be providing other ecosystem services such as comminution of organic matter, nutrient cycling and acting as a food source for other birds and animals (Grodsky et al. 2018). Although vegetation communities were different between reclaimed sites and reference areas, with reference areas exhibiting vegetation communities closer to climax stages of succession, it is important to understand ecosystem dynamics on early seral reclamation sites because land restoration in arid environments is challenging (Shackelford et al. 2021) and sagebrush reestablishment on disturbed sites may take decades (Monroe et al. 2020).

While there was significantly more Rocky Mountain bee plant percent cover on late season well pads than there was overall flower cover on early season well pads, no studies have set a minimum threshold for minimum percent cover of a flowering species to constitute ‘mass flowering.’ As there were no forbs found in reference sites in the early season vegetation monitoring, we assume the restoration areas containing flowers are positively drawing in insects based on research suggesting flower color, scent and height all play roles in being insect attractants (e.g., Wenninger and Inouye 2008). This is the first study to compare insects at more than one point in a growing season on well pads undergoing restoration activities while flowering events are occurring with distinctly different vegetation species. While a total of nine out of 24 insect families found across the two studies were common to both collections, we were unable to determine if insects from early sites traveled to later sites. As insect dispersal studies are becoming more common and affordable (Hagler & Jackson 2001), future research into this area may be beneficial to overall conservation and restoration efforts for insects in oil and natural gas fields. Additionally, determining if insects, which are attracted to these reclamation efforts, are acting as food sources to vertebrate species or providing other ecosystem services to the reference community would be beneficial. This could potentially result in well pads acting as positive reserves instead of negative fragments, making them critical to insect conservation (Cane 2001). Finally, increasing the amount of native forb seeds which can be bought commercially may help improve restoration projects, both by having new plant species increasing ground cover on disturbed sites, and by helping conserve or restore insect habitat and populations and allowing them to continue to provide critical ecosystem services (Curran et al. 2015, Cane and Love 2016, Dumroese et al. 2016).

## Acknowledgements

This study was funded by Jonah Energy and the Wyoming Reclamation and Restoration Center. Dr. Douglas Smith, Department of Ecosystem Science and Management at University of Wyoming, helped with insect identification.

## Notes

### Competing Interest Statement

The authors have declared no competing interest.

## References

Archer, S. and D.A. Pyke. 1991. Plant-animal interactions affecting plant establishment and persistence on revegetated rangeland. Rangeland Ecology and Management 44(6): 558–565.

Belovsky, G.E., and Slade, J.B. 2000. Insect herbivory accelerates nutrient cycling and increases plant production. Proceedings of the National Academy of Sciences, 97(26): 14412–14417.

Cane, J.H. 2001. Habitat fragmentation and native bees: A premature verdict? Ecology and Society 5(1): 3.

Cane, J.H. 2008. Breeding biologies, seed production and species-rich bee guilds of Cleome lutea and Cleome serrulata (Cleomaceae). Plant Species Biology 23 (3): 152–158.

Cane, J.H. and Love, B. 2016. Floral guilds of bees in sagebrush steppe: comparing bee usage of wildflowers available for postfire restoration. Natural Areas Journal 36(4): 377–391.

Curran, M.F., Wolff, B.J. and Stahl, P.D., 2013. Approaching oil and gas pad reclamation with data management: A framework for the future. Journal American Society of Mining and Reclamation, 2(2):195–204.

Curran, M.F., Crow, T.M., Hufford, K.M. and Stahl, P.D., 2015. Forbs and greater sage-grouse habitat restoration efforts: Suggestions for improving commercial seed availability and restoration practices. Rangelands, 37(6): 211–216.

Curran, M.F. and P.D. Stahl. 2015. Database management for large scale reclamation projects in Wyoming: developing better data acquisition, monitoring and models for future applications. Journal of Environmental Solutions for Oil, Gas and Mining, 1(2): 31–43.

Curran, M.F., S.E. Cox, T.J. Robinson, B.L. Robertson, K.J. Rogers, Z.A. Sherman, T.A. Adams, C.F. Strom, and P.D. Stahl. 2019. Spatially balanced sampling and ground level imagery for monitoring reclaimed well pads. Restoration Ecology, 27(5): 974–980.

Curran, M.F., Sorenson, J. and Stahl, P.D., 2019. Rocky mountain beeplant aids revegetation in an arid natural gas field. Environmental Connection, 14: 27–29.

Curran, M.F., Cox, S.E., Robinson, T.J., Robertson, B.L., Strom, C.F. and Stahl, P.D., 2020. Combining spatially balanced sampling, route optimisation and remote sensing to assess biodiversity response to reclamation practices on semi-arid well pads. Biodiversity, 21(4): 171–181.

Curran, M.F., Robinson, T.J., Guernsey, P., Sorenson, J., Crow, T.M., Smith, D.I. and Stahl, P.D., 2022. Insect Abundance and Diversity Respond Favorably to Vegetation Communities on Interim Reclamation Sites in a Semi-Arid Natural Gas Field. Land, 11(4): 527.

Cusser, S., and Goodell, K. 2013. Diversity and distribution of floral resources influence the restoration of plant-pollinator networks on a reclaimed strip mine. Restoration Ecology, 21(6): 713–721.

Dixon, K.W., 2009. Pollination and restoration. Science, 325(5940): 571–573.

Dumroese, R.K., T. Luna, J.R. Pinto, and T.D. Landis. 2016. Forbs: Foundation for restoration of monarch butterflies, other pollinators, and greater sage-grouse. Natural Areas Journal 36(4): 499–511.

Grodsky, S.M., R.B. Iglay, C.E. Sorenson, C.E. Moorman. 2018. Should invertebrates receive greater inclusion in wildlife research journals? The Journal of Wildlife Management 79(4): 529–536.

Hagler, J.R., and Jackson, C.G. 2001. Methods for marking insects: current techniques and future prospects. Annual Review of Entomology, 46(1): 511–543.

Harmon, J.P.; Ganguli, A.C.; Solga, M.A. 2011. An overview of pollination in rangelands: who, why, and how. Rangelands, 33(3): 4–9.

Holzschuh, A., Dormann, C. F., Tscharntke, T., & Steffan-Dewenter, I. 2013. Mass-flowering crops enhance wild bee abundance. Oecologia, 172(2): 477–484.

Lavelle, P. 1997. Soil function in a changing world: the role of invertebrate ecosystem engineers. European Journal of Soil Biology, 33:159–193.

Lockwood, J. 2013. The infested mind: Why humans fear, loathe, and love insects. Oxford University Press, Oxford, UK.

Longcore, T. 2003. Terrestrial arthropods as indicators of ecological restoration success in coastal sage scrub (California, USA). Restoration Ecology 11(4): 397–409.

Losey, J.E., and M. Vaughan. 2006. The economic value of ecological services provided by insects. Bioscience 56(4): 311–323.

Menz, M.M.H., Phillips; R.D., Winfree, R., Kremen, C., Aizen, M.A., Johnson, S.D. and Dixon, K.W. 2011. Reconnecting plants and pollinators: challenges in the restoration of pollination mutualisms. Trends in Plant Sciences, 16(1): 4–12.

Milne, L.J. and Milne, M. 2011. National Audubon Society Field Guide to Insects and Spiders: North America. Toppon Printing Co., Tokyo, Japan.

Monroe, A.P., Aldridge, C.L., O’Donnell, M.S., Manier, D.J., Homer, C.G. and Anderson, P.J., 2020. Using remote sensing products to predict recovery of vegetation across space and time following energy development. Ecological Indicators, 110: 105872.

Noriega, J.A., Hortal, J., Azcárate, F.M., Berg, M.P., Bonada, N., Briones, M.J., Del Toro, I., Goulson, D., Ibanez, S., Landis, D.A. and Moretti, M. 2018. Research trends in ecosystem services provided by insects. Basic and Applied Ecology, 26: 8–23.

Rohde, A.T., Pilliod, D.S. and Novak, S.J. 2019. Insect communities in big sagebrush habitat are altered by wildfire and post-fire restoration seeding. Insect Conservation and Diversity, 12(3: 216–230.

Samways, M.J., Barton, P.S., Birkhofer, K., Chichorro, F., Deacon, C., Fartmann, T., Fukushima, C.S., Gaigher, R., Habel, J.C., Hallmann, C.A. and Hill, M.J., 2020. Solutions for humanity on how to conserve insects. Biological Conservation, 242: 108427.

Shackelford, N., Paterno, G.B., Winkler, D.E., Erickson, T.E., Leger, E.A., Svejcar, L.N., Breed, M.F., Faist, A.M., Harrison, P.A., Curran, M.F. and Guo, Q. 2021. Drivers of seedling establishment success in dryland restoration efforts. Nature Ecology & Evolution, 5(9): 1283–1290.

Shelomi, M. 2015. Why we still don’t eat insects: Assessing entomophagy promotion through a diffusion of innovations framework. Trends in Food Science & Technology, 45(2): 311–318.

Stahl, P.D. and Curran, M.F., 2017. Collaborative efforts towards ecological habitat restoration of a threatened species, Greater Sage-Grouse, in Wyoming, USA. In Land Reclamation in Ecological Fragile Areas (pp. 251–254). CRC Press.

Sylvain, Z.A., E.K. Espeland, T.A. Rand, N.M. West, and D.H. Branson. 2019. Oilfield reclamation recovers productivity but not composition of arthropod herbivores and predators. Environmental Entomology, 48(2): 299–308.

Tallamy, D.W. 2004. Do alien plants reduce insect biomass? Conservation Biology 18(6): 1689–1692.

Tallamy, D. 2009. Bringing Nature Home: How Native Plants Sustain Wildlife in our Gardens, updated and expanded. Timber Press, Portland, OR, USA.

Tonietto, R.K. and Larkin, D.J. 2018. Habitat restoration benefits wild bees: A meta-analysis. Journal of Applied Ecology, 55(2): 582–590.

Todd, K.J., M.M. Gardiner, and E.D. Lindquist. 2016. Mass flowering crops as a conservation resource for wild pollinators (Hymenoptera: Apoidea). Journal of the Kansas Entomological Society, 89(2): 158–167.

Wenninger, E.J., and Inouye, R.S. 2008. Insect community response to plant diversity and productivity in a sagebrush-steppe ecosystem. Journal of Arid Environments, 72(1): 24–33.

Westphal, C.; Steffan-Dewenter, I.; Tscharntke, T. 2003. Mass flowering crops enhance pollinator densities at landscape scales. Ecology Letters, 6(11): 961–965.

