## Supplemental Table 1 for "Early seral vegetation communities increase insect abundance and diversity in a semiarid natural gas field during early and late growing season"

**Species in Seed Mix 'B1' in Jonah Field**

*Achnatherum hymenoides*
*Elymus elymoides* (Raf.) Swezey spp. *brevifolius*
*Pseudoroegneria spicata* (Pursh)A. Love
*Leymus cinereus* (Scribn. & Merr.) A. Love
*Hesperostipa comata* (Trin. & Rupr.) Barkworth
*Atriplex canescens* (Pursh) Nutt.
*Sphaeralcea munroana* (Douglas) Spach
*Oenothera pallida* Lindl.
*Penstemon procerus* Douglas ex Graham
*Cleome serrulata* Pursh
*Penstemon palmeri*
*Artemisia tridentata* spp. *Wyomingensis* Beetle & Young
*Achillea millefolium* L. var. *occidentalis* DC.
*Eriogonum umbellatum* Torr.
*Artemisia tridentata* Nuttall
*Symphyotrichum laeve* (L.) A. Love & D. Love var. *leave*
*Lupinus argenteus* Pursh
*Linum lewisii* Pursh
*Krascheninnikovia lanata* (Pursh) A. Meesuse & Smit
*Poa secunda* Presl
*Erigeron engelmannii* A. Nelson
